# Convergent clonal selection of donor- and recipient-derived CMV-specific T cells in hematopoietic stem cell transplant patients

**DOI:** 10.1101/2021.09.14.458942

**Authors:** Jami R. Erickson, Terry Stevens-Ayers, Florian Mair, Bradley Edmison, Michael Boeckh, Philip Bradley, Martin Prlic

**Author notes:** Co-Corresponding Authors: Martin Prlic, 1100 Fairview Ave N, Seattle, WA 98109, Philip Bradley, 1100 Fairview Ave N, Seattle, WA 98109.

## Abstract

Competition between antigen-specific T cells for peptide:MHC (p:MHC) complexes shapes the ensuing T cell response. Mouse model studies provided compelling evidence that competition is a highly effective mechanism controlling the activation of naïve T cells. However, assessing the effect of T cell competition in the context of a human infection requires defined pathogen kinetics and trackable naïve and memory T cell populations of defined specificity. A unique cohort of non-myeloablative hematopoietic stem cell transplant (nmHSCT) patients allowed us to assess T cell competition in response to CMV reactivation, which was documented with detailed virology data. In our cohort, HSCT donors and recipients were CMV-seronegative and -positive, respectively, thus providing genetically distinct memory and naïve T cell populations. We used single-cell transcriptomics to track donor versus recipient-derived T cell clones over the course of 90 days. We found that donor-derived T cell clones proliferated and expanded substantially following CMV-reactivation. However, for immunodominant CMV epitopes, recipient-derived memory T cells remained the overall dominant population. This dominance was maintained despite more robust clonal expansion of donor-derived T cells in response to CMV reactivation. Interestingly, the donor-derived T cells that were recruited into these immunodominant memory populations shared strikingly similar TCR properties with the recipient-derived memory T cells. This selective recruitment of identical and nearly identical clones from the naïve into the immunodominant memory T cell pool suggests that competition does not interfere with rejuvenating a memory T cell population, but results in selection of convergent clones to the memory T cell pool.

**Significance:** An existing memory T cell population specific for a single epitope is sufficient to effectively curtail responses to any new antigens if the original epitope is present in a vaccination regimen or heterologous infections. We asked if T cell competition precludes recruitment of any new, naïve T cells to an existing memory T cell pool in context of CMV-specific T cell responses in a cohort of transplant patients. Our data indicate that competition does not prevent recruitment of naïve T cells into the memory T cell pool, but selects for T cells with nearly or fully congruent T cell receptor specificities. We discuss the implications of rejuvenating a memory T cell pool while preserving the T cell receptor repertoire.

## Introduction

The human T cell compartment is estimated to contain 10^12^ cells (1). This T cell compartment consists of clonally expanded memory T cells and a pool of naïve T cells with an estimated ~10^8^ unique TCRβ chains (1). Competition between different T cell populations for resources and niches has been studied for nearly 50 years (2). With the development of the p:MHC tetramer and TCR transgenic mouse models, assessing competition between different T cell populations for a given p:MHC epitope has become more nuanced in the past 20 years (3–5). Initial observations focused on T cell responses against alloantigens in a transplant context (2, 6, 7), while subsequent studies examined competing T cell responses during infections or mimicking infection-like scenarios by using peptide-pulsed antigen-presenting cells (APCs) (8, 9). T cell fitness and competition are shaped by TCR affinity for p:MHC, the T cell precursor frequency, and even epitope-independent cross-competition (3, 10, 11). In cross-competition, T cells compete for access to APCs instead of just their specific p:MHC (3, 9). Cross-competition does not appear to occur in all infections (12), but the mechanisms that control the extent of cross-competition remain poorly defined. Of note, a memory T cell population specific for just one single epitope can very effectively limit *de novo* T cell responses to other epitopes present in a subsequent heterologous infection or vaccine boost scenario as reported using different mouse model systems (4, 5). Similar findings were reported in a human study with a cohort that was suitable to examine the effects of T cell competition (13). In this latter study, Frahm and colleagues found that pre-existing T cell memory to adenovirus serotype 5 (Ad5) could substantially limit the response to HIV-derived epitopes delivered by an Ad5-vectored HIV vaccine (13).

Addressing T cell competition in human cohorts is challenging as it requires distinction between memory T cell responses versus *de novo* responses in context of a well-defined priming event. Furthermore, it requires strong T cell responses so analysis of a limited number of T cells from the peripheral blood is sufficient to detect antigen-specific T cell clones. One of the strongest and best characterized human T cell responses occurs in response to cytomegalovirus (CMV) infection. Given that T cell responses to the immunodominant CMV proteins pp65 and IE1 can be found in most CMV-seropositive individuals (14–19), we reasoned that assessing T cell competition for pp65- and IE1-derived epitopes could be feasible. To study competition, we examined cryopreserved peripheral blood mononuclear cells from patients undergoing a non-myeloablative hematopoietic stem cell transplant (nmHSCT) with concomitant monitoring for CMV reactivation. Importantly, an nmHSCT preserves a substantial amount of the patient’s immune system compared to conventional HSCTs (20–22). Although CMV-seropositive individuals are highly prevalent (23), we identified a small cohort of nmHSCT patients with CMV-seronegative donors. Given that recipient and donor cells are genetically distinct, samples from this cohort represent a unique opportunity to measure competition between recipient-derived CMV-specific memory T cells and donor-derived naïve T cells. To simultaneously distinguish donor- and recipient-derived cells, and assess T cell repertoire and specificity, we utilized single-cell transcriptomics, ex vivo stimulation assays and computational analysis approaches. Overall, we found that T cells specific for CMV immunodominant epitopes remained predominantly recipient-derived, while clonal expansion following CMV reactivation was mainly driven by donor-derived T cells. We also observed that donor-derived T cells were not excluded from entering the immunodominant (pp65 and IE-1) CMV-specific memory CD8+ T cell pool. Finally, some donor-derived T cell clones showed stunning similarities in TCR α and β V(D)J usage including highly conserved (and in one instance even identical at the amino acid level) CDR3 regions with the recipient-derived memory T cell pool. Overall, our data suggest that the recipient-derived, CMV-specific memory T cell pool is quickly rejuvenated by newly recruited donor-derived clones. We discuss basic immunology implications as well as clinical relevance of our findings.

## Results

### Patient cohort, sample processing and virology data

To study competition between memory and naïve T cells for the same antigen, we obtained longitudinal samples from patients who underwent nmHSCT (**Figure 1A**). The patients were selected based on the following criteria: 1) recipients were CMV seropositive; 2) donors were CMV seronegative, allowing us to track the naïve T cell response to CMV; 3) both donors and recipients expressed the HLA-A*02:01 allele, allowing us to identify CMV-specific CD8+ T cells using published TCR sequences and a p:MHC tetramer (epitope NLVPMVATV from CMV); and 4) CMV reactivation occurred between 30 and 90 days after transplant. Following nmHSCT, each of the four patients underwent regular peripheral blood draws and weekly CMV surveillance by PCR (24). Blood draws were used to assess white blood cell (WBC) counts and processed to cryopreserve peripheral blood mononuclear cells (PBMCs) (**Figure 1B**). Cryopreserved PBMCs from each patient at three different time points - days 30, 60 and 90 post nmHSCT (red lines indicate PBMC draws that were used for each patient, **Figure 1C**) - were run through our single cell analysis pipeline. This pipeline included assays to determine cellular phenotypes, transcriptional profiles, T cell receptor (TCR) sequences and TCR specificities. We utilized transcriptional and machine learning analysis, such as TCRDist (25), to determine how clonal populations respond to CMV reactivation over time and predict shared epitope specificity of distinct clones. Finally, detailed virologic data were collected for each patient (**Figure 1C**). Patients 1, 2 and 4 had CMV reactivation events between days 60 and 90. Patient 3 had a CMV reactivation event between days 30 and 60 (**Figure 1C**).

**Figure 1:**
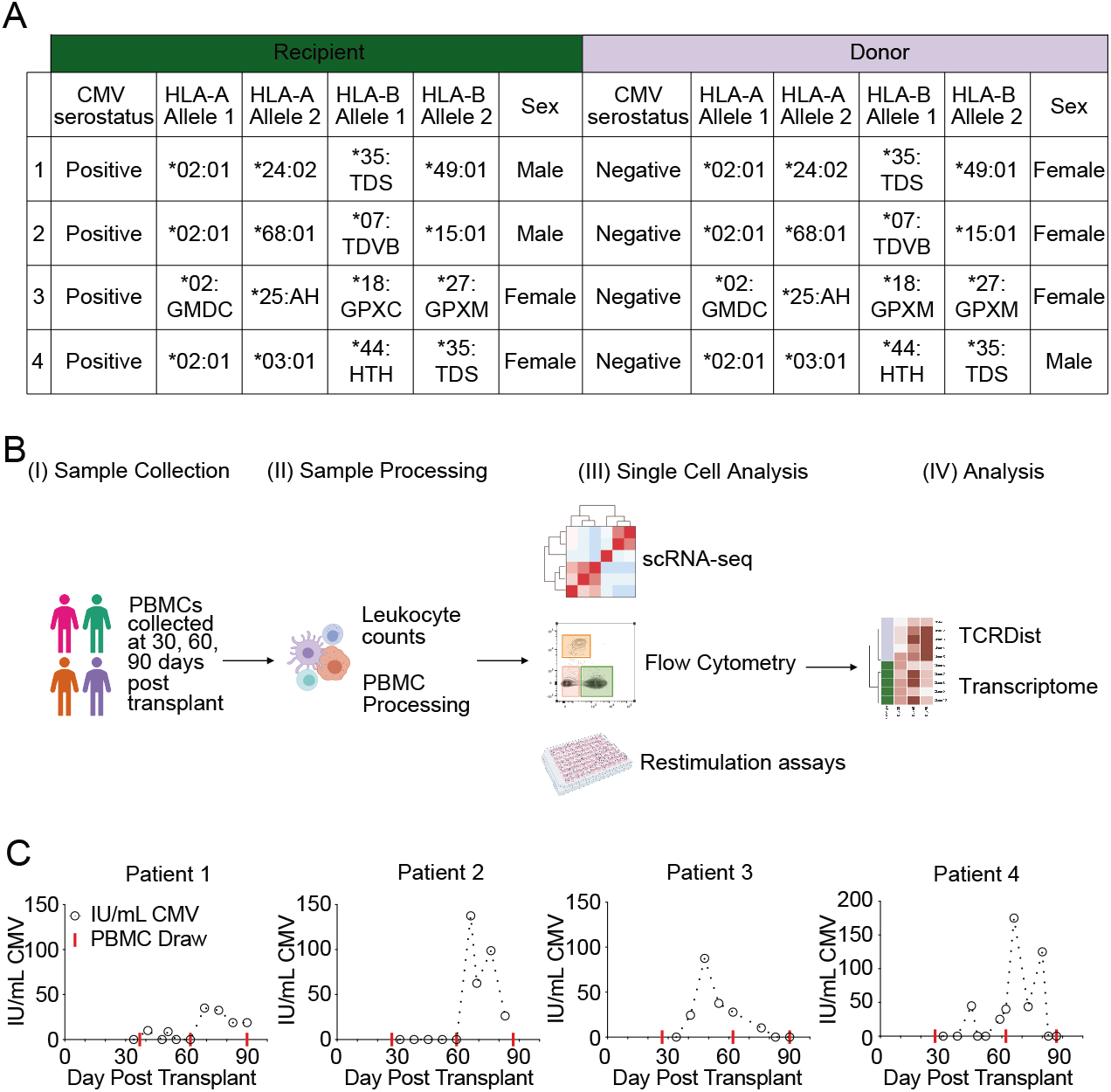
Overview of the patient cohort, the experimental design and collected virology data. (A) Patient cohort details for each recipient and donor pair. (B) Overview of our experimental strategy. PBMCs: Peripheral blood mononuclear cells. scRNA-seq: single-cell RNA-sequencing. (C) Cytomegalovirus (CMV) load over time for each patient. IU/mL: international units per mL of CMV. Red line indicates blood draws that scRNA-seq was performed.

### Immune compartment composition of the nmHSCT cohort on days 30, 60 and 90 post transplant

First, we established baseline values of the immune compartment in our patient cohort by evaluating the nmHSCT recipients prior to CMV reactivation. Using hematology reports and flow cytometry data, we calculated absolute numbers of white blood cells, T cells, and CD8+ T cells for each patient (**Figure 2A**). In three of the four patients, the number of both T cells and CD8+ T cells increased over time. While patient 2 had decreasing absolute numbers of T cells over time, all patients had an increase in the percentage of T cells (**Figure 2B**).

**Figure 2:**
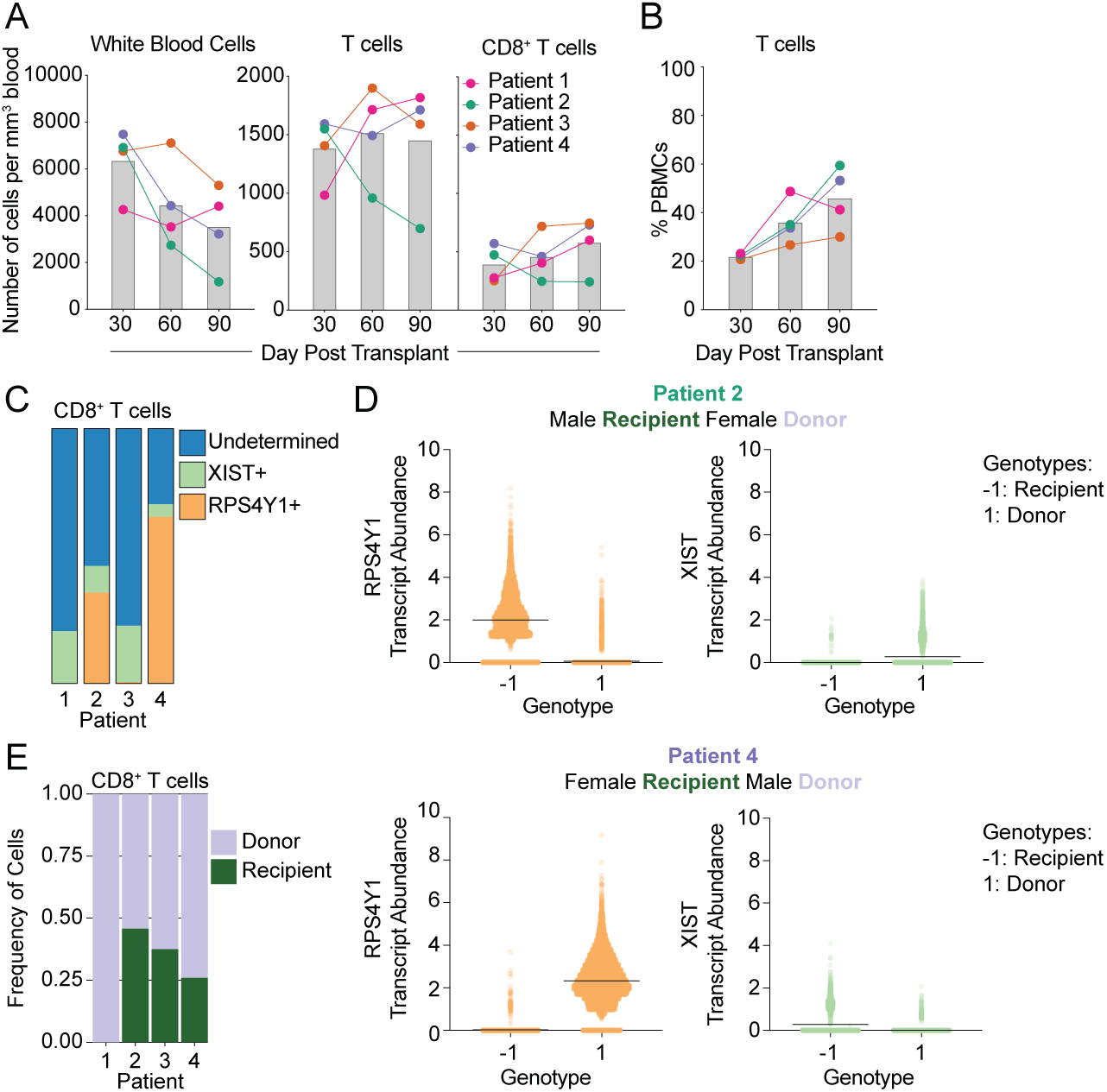
Immune compartment composition of the nmHSCT cohort on days 30, 60 and 90 post transplant. (A) Absolute number of leukocytes, T cells, and CD8^+^ T cells per mm^3^ for each patient over time. Grey bars represent the average of all patients. (B) Percentage of T cells of live PBMCs over time for each patient. Grey bars represent the average of all patients. (C) Stacked-bar plot representing the number of CD8+ T cells that are RPS4Y1+, XIST+, or undetermined (either RPS4Y1-XIST- or RPS4Y1+ XIST+) in each patient. (D) Each cell was called for a genotype (either −1 or 1) based on single nucleotide polymorphisms. Then sex-linked gene expression was assessed for each genotype to determine which genotype belonged to recipient and donor. (E) Stacked-bar plot of CD8+ T cells for each patient displaying the frequency of recipient and donor cells.

In order to assess competition between memory and naïve T cells for the same p:MHC, we needed to delineate recipient cells from donor cells in the nmHSCT patients. First, we attempted to do this by examining sex-linked gene expression from our single-cell RNA sequencing (scRNA-seq) data as three out of four of the patients (1, 2, and 4) received sex-mismatched transplants (**Figure 1A**). In particular, we utilized expression of two genes: the male specific *RPS4Y1* (a Y chromosome-linked ribosomal protein) and the female specific *XIST* (a long non-coding RNA used for X chromosome inactivation). Both *RPS4Y1* and *XIST* transcripts could be found in Patient 2 and 4, allowing us to use sex-mismatched transplants to differentiate donor from recipient in these patients (**Figure 2C**). However, due to low detection and high dropout rates inherent in scRNA-seq (26–28), the vast majority of cells were of undetermined origin (**Figure 2C**, blue bars). Of note, patient 1 was male, but had undergone a previous HSCT procedure and we could not detect any RPS4Y1-expressing (recipient-derived) T cells in this patient. Further, patient 3 of our cohort had a sex-matched donor and recipient. In an effort to improve donor vs. recipient identification, we used single nucleotide polymorphisms (SNPs) within the scRNA-seq data to differentiate donor from recipient cells. SNP analysis had already been conducted on patient 3 (29), which allowed us to match SNPs in our scRNA-seq data to these previously acquired data and define donor vs. recipient. To resolve recipient and donor within patients 2 and 4, we utilized the scRNA-seq data to find SNPs (N=823 and 700 for patients 2 and 4, respectively) within the transcripts for each cell. We employed these identified SNP markers to assign each cell one of two different genotypes (−1 or 1). Once each cell was assigned a genotype, we then designated recipient or donor by using sex-linked gene expression for each genotype (**Figure 2D**). Together, this allowed us to observe the contribution of recipient and donor to the CD8+ T cell compartment (**Figure 2E**). Overall, each patient contained proportionally more donor than recipient-derived CD8+ T cells (**Figure 2E**).

### Assessing the presence of donor and recipient-derived pp65:A02:01-specific T cells

As a first step, we wanted to ensure that we could indeed detect CMV-specific CD8+ T cell responses in all 4 patients. We used a p:MHC tetramer to identify CD8+ T cells that were specific for the immunodominant pp65 peptide NLVPMVATV presented in the context of HLA-A02:01 for all 4 patients. We detected pp65:A02:01-specific T cells in all 4 patients across all 3 time points (**Fig. 3A**). We next asked whether any of these pp65:A02:01-specific T cells were donor-derived by day 90. To address this, we sorted pp65:A02:01-specific T cells and then analyzed the sorted cells using 3’ scRNA-seq to determine whether each cell originated from the donor or recipient as described above. We found that 90 days after nmHSCT, the vast majority of pp65:A02:01-specific T cells were still recipient-derived in patients 2 and 3 (**Fig. 3B**), despite the majority of the immune compartment being donor-derived. Of note, patient 1 had more pp65:A02:01+ T cells that were donor-derived, but since this patient had undergone a transplant procedure prior to the nmHSCT, the vast majority of the immune compartment was donor-derived (**Fig. 2E**). Due to a technical issue, we only interrogated a fairly limited population of pp65:A02:01+ T cells of Patient 4 in our scRNA-seq analysis (**Fig. 3B**).

**Figure 3:**
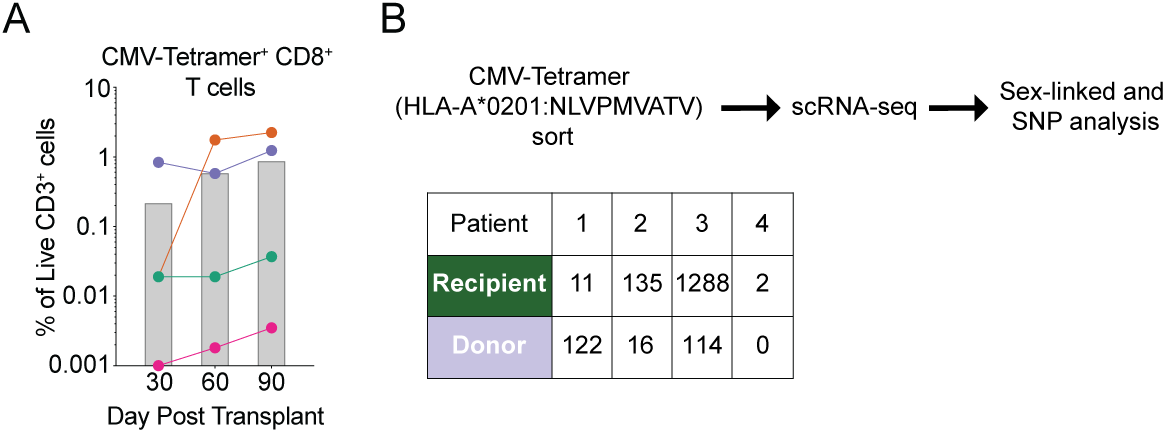
CMV-specific T cells are present in all 4 patients. (A) Frequency of HLA-A2 NLV tetramer-specific T cells in each patient on days 30, 60 and 90. Grey bars represent the average of all patients. (B) Overview of workflow to determine CMV-tetramer+ CD8+ T cell origin. (C) A table indicating the number of HLA-A2 NLV tetramer-specific CD8+ T cells derived from recipient or donor for patients 2-4 at 90 days post-transplant.

Overall, these data indicate that the cohort is suitable to examine T cell competition and also provide a first line of evidence that although T cell competition appears to occur, it is permissive and allows for the recruitment of new T cell clones from the donor-derived, antigen-naïve T cell population.

### Assess clonal expansion in the CD8+ T cell compartment

After establishing that the cohort is suitable to address competition, we first wanted to determine how many expanding T cell clones we could detect in the entire CD8 T cell compartment. We reasoned that assessing expanding TCR clones would provide a general overview of the CD8+ T cell dynamics, which was needed as a reference to subsequently assess and interpret CMV-specific T cell responses.

To accomplish this, we performed 5’ scRNA-seq on CD8+ T cells to acquire VDJ and transcriptome data, followed by identifying TCR clones that expand over time (**Fig. 4A**). Note that this CD8 T cell population did not contain any pp65:A02:01 tetramer+ T cells, since these cells were sorted and analyzed separately (**Fig. 3B**). We defined “expanding clones” as having 2-fold more cells than the previous timepoint and having at least 10 cells at any time point. We observed that the majority of these expanding clones were donor-derived (**Fig. 4A**). We next quantified the fold change in frequency of these expanding clones between day 60 and 90 and we observed that donor-derived expanding clones had an ~10-fold increased rate of fold expansion compared to recipient-derived clones (**Fig. 4B**). Together, these data indicate that clonal expansion is dominated by donor-derived T cells, which also appear to expand more vigorously. We next wanted to estimate the number of cell divisions that occurred within each donor-derived expanding clone. We first determined the relative abundance of each expanding clone within the CD8+ T cell compartment using our scRNA-seq data. We used the flow cytometry data and the absolute WBC numbers from each patient’s hematology reports to extrapolate the absolute numbers of each clone per mm^3^ blood. Assuming an average of 5L of blood per person and assuming each CD8+ T cell clone was only present as a single cell at the time of priming, we found that on average, expanding clones would undergo nearly 24 rounds (±1.56 standard deviation) of cell division (**Fig. 4C**). The number of rounds of cell division was calculated by determining absolute numbers, then determining the number of doubling events that would occur to lead to that number of cells. The clones that were included were considered ‘expanding clones’, and had zero clones detected at day 30 post-transplant. Although this number may be an overestimate if the single clone progenitor assumption is incorrect, it indicates that clonal expansion was robust and did not appear to be stifled by mechanisms related to competition or cross-competition.

**Figure 4:**
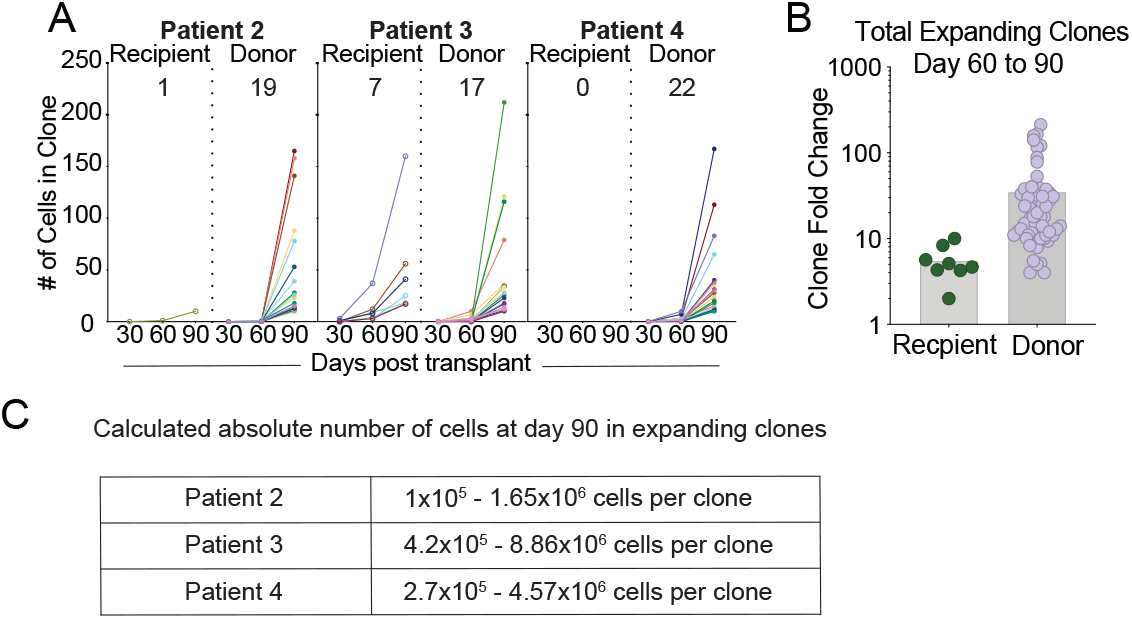
Assessing general patterns of clonal expansion in the CD8+ T cell compartment. (A) Plots indicating the number of cells within a CD8+ T cell clone over time, separated by clone origin. Left to right: (1) All Clones, (2) clones expanding at day 60 post-transplant, (3) clones expanding at day 90 post-transplant, (4) contracting clones, and (5) stable clones. (B) Fold-expansion of expanding donor- and recipient-derived clones is shown between day 60 and 90. (C) A calculated range of abundance for expanding clones in the blood.

### Additional CD8+ T cell clonal analysis reveals the presence of stable, expanding and contracting clones

We next wanted to better define the overall dynamics of CD8+ T cell clone abundance and determine the relative abundance of expanding, stable and contracting clones. First, we examined the trends of all CD8+ T cell clones over time. To do this, we plotted each clone that had at least five cells per clone for each time point (**Fig. 5A**). We then separated each of the clones by behavior, with four different behaviors: 1) clones that expanded at 60 days post-transplant, 2) clones that expanded at 90 days post-transplant, 3) clones that contracted, and 4) clones that remained stable over time. Our criteria for clones that expanded at 60 days post-transplant were that the clone must have had at least five cells at one time point and the number of cells at day 60 had to be twice the number of cells at day 30. We found that each patient had clones that expanded at day 60, but the majority of the clones which expanded at day 60 were recipient derived clones from patient 3. In contrast, the majority of clones which expanded at day 90 were donor-derived. Contracting clones were defined as those clones which decreased two-fold between any two time points and had at least 5 cells at any time point. Clones that were considered stable had at least 5 cells at any time point and did not vary by more than two-fold at any time point.

**Figure 5.**
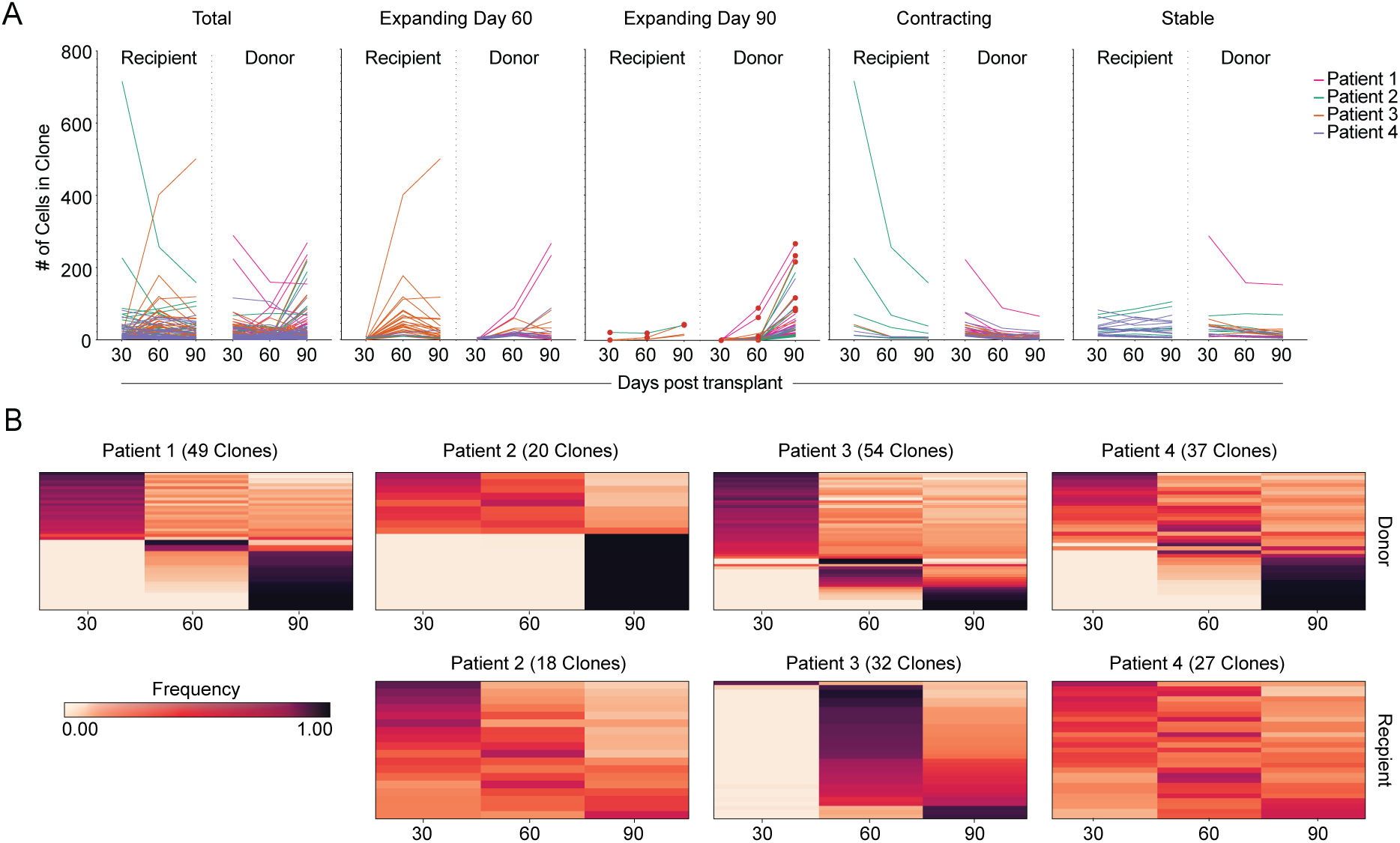
CD8 T cell compartment expansion and contraction dynamics. (A) The number of total detected clones is shown (left panel) and then divided into expanding (day 30 to 60; day 60 to 90), contracting and stable clones. (B) Frequency heat maps of clones separated by patient and clone origin. Each row represents one clone. The color represents the frequency of that clone at each time point post-transplant.

To further illustrate how the frequencies of each clone changed over time, we used frequency maps of the clones from each patient with donor-derived clones shown in the top row and recipient-derived clones in the bottom row (**Fig. 5B**). We found that all patients contained at least some donor-derived clones that reached peak frequency at 90 days post-nmHSCT. In contrast, most recipient-derived clones remained stable or contracted over time, although some recipient-derived clones from patient 3 reached peak frequency at either 60- or 90-days post-nmHSCT, respectively. Overall, these data further highlight that clonal expansion is predominantly observed in the donor-derived T cell population.

### Expanding clones have similar transcriptional profiles regardless of donor or recipient origin

Although most expanding clones were of donor origin, some expanding T cells were recipient-derived clones. We next wanted to assess if donor- and recipient-derived expanding T cells had congruent or distinct transcriptional programs. We considered that transcriptional differences could result from distinct epitope specificity, distinct cell origin (donor vs. recipient) and differentiation status at the time of activation (naïve vs. memory). We examined how transcriptional profiles change over time by focusing on clones (either donor-derived, or recipient-derived) that expanded at 90 days post-nmHSCT. To visualize gene expression changes for these single cell data, we used uniform manifold approximation and projection (UMAP) (30). CD8+ T cells were then placed into clusters identified by a shared nearest neighbor modularity optimization-based clustering algorithm. Each clone was visualized on the CD8+ T cell UMAP at each time point, and each cell within that clone was colored by cluster (**Fig. 6A**). Note, that for patients 1 and 4 all plots shown are of donor origin. In general, the clones which expanded at 90 days post-nmHSCT occupied the same five clusters in the UMAP, which are displayed as green, blue, brown, purple and red. This pattern could be observed regardless if clones were donor- or recipient-derived indicating a shared transcriptional phenotype. We used singleR to perform cell type calling (31), which suggested that cells occupying these clusters featured an effector memory CD8+ T cell phenotype. There was a unique phenotype seen in some recipient-derived clones of patient 3 that occupied an “orange” cluster that fell into a distinct spatial region and contains a transcriptome suggestive of a terminal effector phenotype (based on singleR designation). Overall, these data suggest that T cell clones which expanded at 90 days post-nmHSCT had a similar transcriptional profile, regardless of epitope specificity or donor vs. recipient origin. Of note, these expanding clones share the same transcriptional space even across patients. The main characteristics of this transcriptional profile as characterized by gene ontology analysis include “effector molecules” (including *GZMH*, *GZMB*, *IFNG*), “TCR mediated signaling” (including *CD3D*, *LCK*, *LAG3*), and “memory T cell formation” (including *ZEB2*, *CCL5*, *LGALS1*).

**Figure 6.**
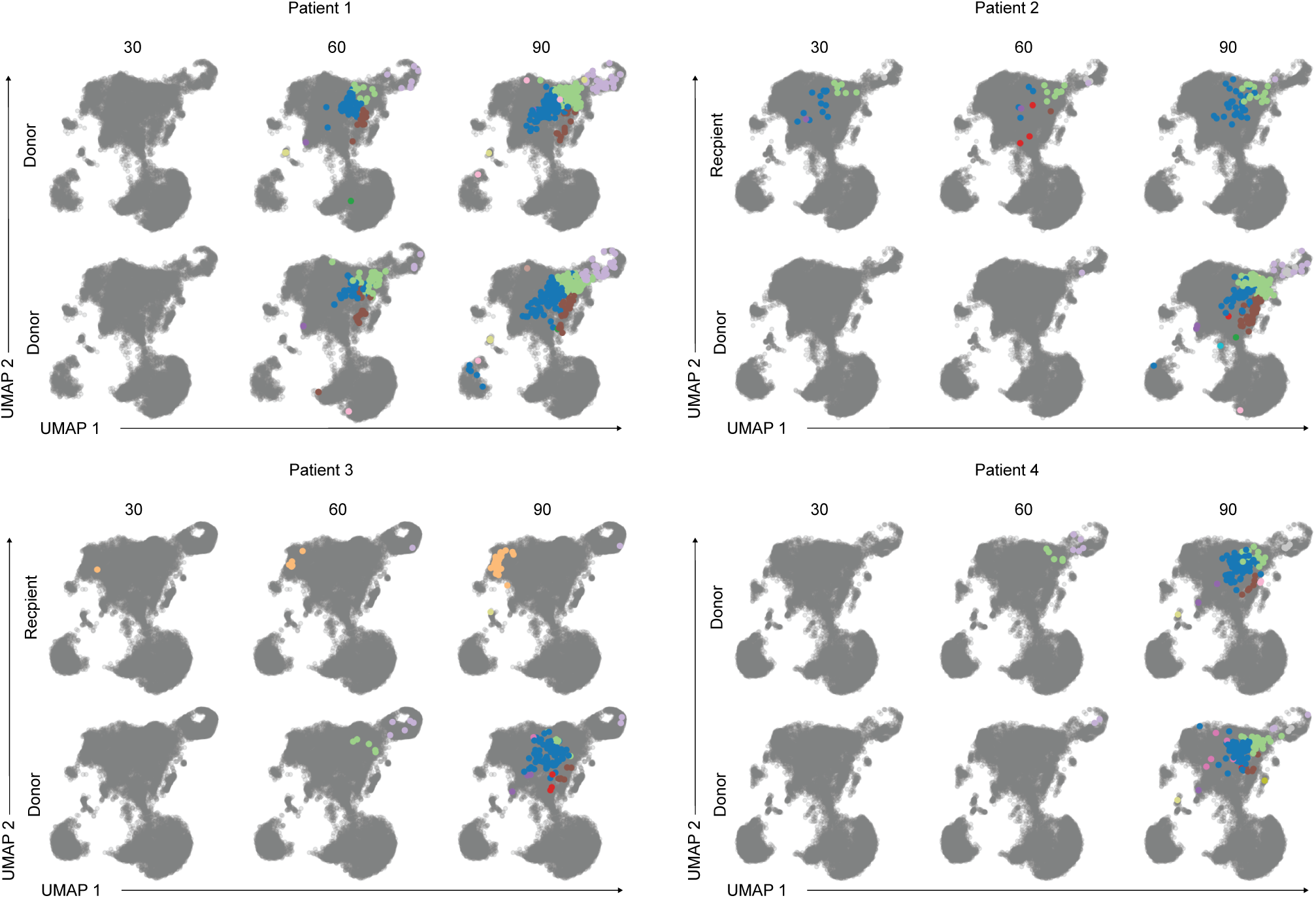
Shared transcriptional signatures of expanding CD8 T cell clones. (A) UMAPs of expanding clones from each patient. The color of the cell represents cluster identity. The grey UMAP indicates where other cells from each patient reside.

### Exploring CMV-specific T cell responses

We next wanted to examine if we could identify CMV-specific T cells within these expanding clones. To achieve this, we needed to expand our analysis to T cells beyond those that we already identified by pp65 peptide (NLVPMVATV) loaded HLA-A*02:01 tetramers. As an overall strategy, we used a combination of searching for previously published sequences, identifying CMV-specific T cells in ex vivo stimulation experiments and using TCRdist analysis to assign CMV-specificity to TCR clones.

First, we compared the V(D)J full chain sequences of clones from our cohort to previously published V(D)J sequences of CMV-specific CD8+ T cells (32). We identified 15 clones in our cohort with sequences that had been previously described as being CMV-specific (9 recipient-derived, 6 donor-derived) (**Fig. 7A**).

**Figure 7.**
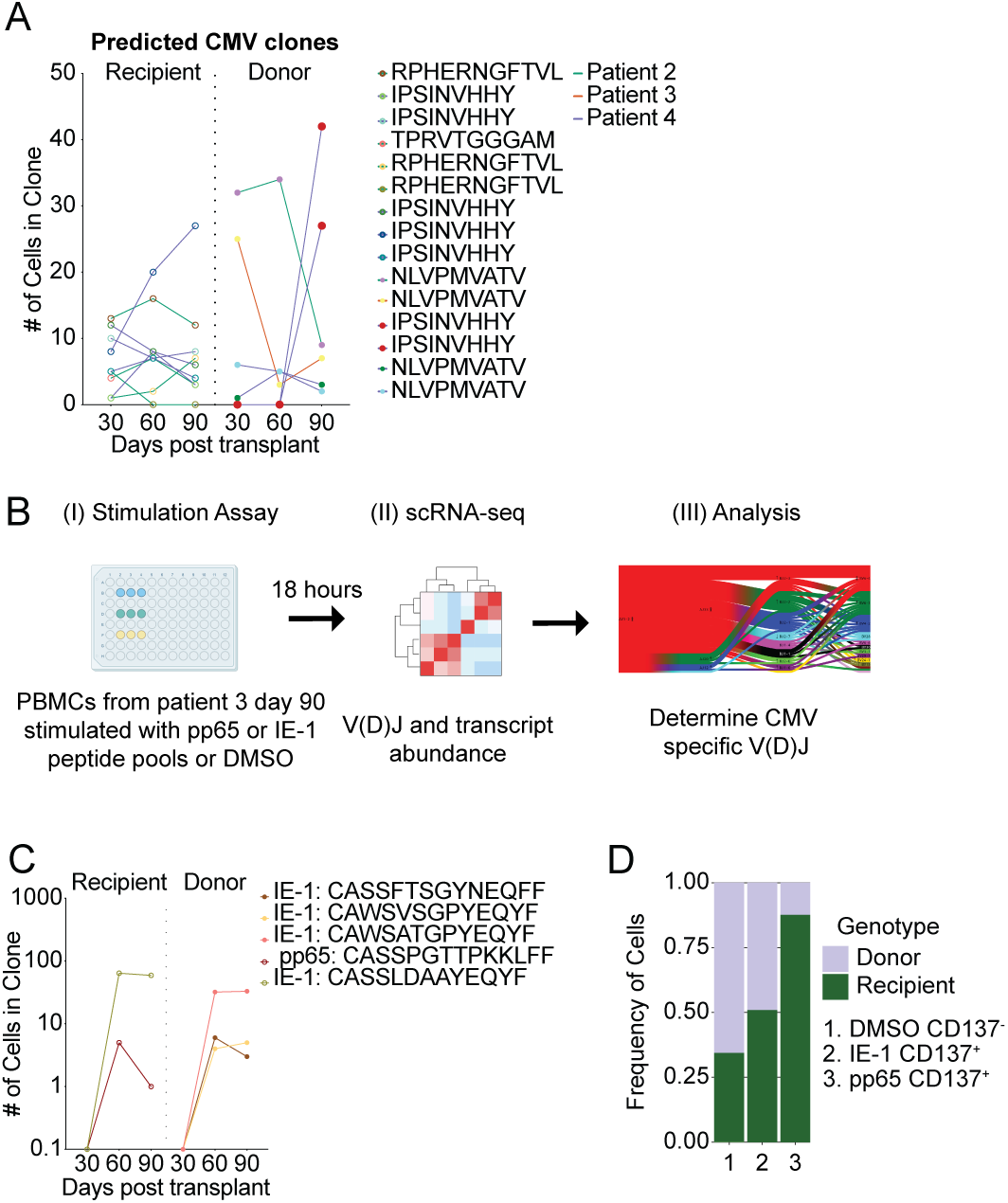
Identification of CMV-specific CD8 T cell clones. (A) CMV-specific CD8 T cell clones were identified based on previously published reports. The number of cells for each CMV-specific CD8 T cell clone is shown over time. Circle color indicates clonal specificity for a given peptide. Line color indicates which patient the clone is from. (B) Experimental overview of stimulation assay. PBMCs from patient 3 at day 90 post-transplant were stimulated with pp65 or IE-1 peptide pools for 18 hours. FACS-purified T cells were analyzed by scRNA-seq followed by V(D)J analysis was performed to determine CMV specific V(D)J sequences. (C) CMV-specific CD8 T cell clones were identified in an ex vivo stimulation assay for patient 3 as shown in (B). The number of cells for each identified clone is plotted over time. Circle color indicates clonal specificity for a given peptide. (D) Frequency of donor and recipient cells is shown for the negative control (DMSO), the IE-1 peptide pool stimulation condition and the pp65 peptide pool stimulation condition.

Second, to identify additional CMV-specific T cell receptors, we stimulated T cells from patient 3 using overlapping peptide pools from two CMV proteins: pp65 and IE-1. We chose patient 3 given the high abundance of pp65:A02:01-specific T cells (**Fig. 3A**) and reasoned that this would likely yield the highest number of new CMV-specific TCR specificities. PBMC were incubated with either dimethyl sulfoxide (DMSO, carrier control), pp65 peptide pool, or IE-1 peptide pool for 18 hours (**Fig. 7B**). Following incubation, we used CD137 as a surrogate marker for responding, antigen specific T cells (33). We used flow cytometry to sort the following populations after stimulation: CD137-CD8+ T cells (DMSO), CD137+CD8+ T cells (pp65 peptide pool), and CD137+CD8+ T cells (IE-1 peptide pool). Each sorted population was then analyzed by 5’ scRNA-seq to determine the V(D)J sequences. Of note, CD8+ T cells that were treated with either peptide pool expressed more effector molecule transcripts compared to DMSO treated controls (**Supp. Fig. 1A**). Sequencing of the CD137+CD8+ T cells from the pp65 stimulation conditions identified 27 clones. Similarly, 14 clones were identified by sequencing CD137+CD8+ T cells from the IE-1 peptide pool stimulation. We compared the TCR sequences of these new clones to the original longitudinal patient 3 samples (**Supp. Fig. 1B**). Out of these 41 new clones, we found five clones that were also present in our initially generated *ex vivo* data set (**Fig. 7C**). One of the clones was specific for a pp65-derived epitope and the other four clones were specific for IE-1-derived epitopes. We did not expect a complete overlap between these datasets given that the sampling itself is inherently limited (number of T cells sequenced in each experiment as a snapshot of the entire CD8 repertoire), but we were initially surprised by this rather limited congruence. However, since we did not sequence pp65:A02:01-specific T cells in our initial experiment (since the tetramer+ CD8 T cells were sequenced separately), the low number of pp65:MHC-I-specific clones could indicate that most pp65-specific T cells are truly pp65(NLVPMVATV):A02:01-specific. Finally, of the five pp65- and IE1-specific clones identified, only two clones met our criteria for expanding clones, and both of those clones expanded from day 30 to day 60 post-transplant, which aligns with the CMV reactivation kinetics for donor 3 (between day 30 and 60).

We next determined recipient and donor contribution of both pp65:MHC-I and IE-1:MHC-I specific CD8+ T cells. CD137-CD8+ T cells (DMSO control) were mostly donor derived (723 recipient cells and 1375 donor cells, **Fig. 7C**), and yielded a comparable donor to recipient distribution as initially observed in **Fig. 2E**. However, pp65 or IE-1-specific T cells had an increased frequency of recipient cells similar to our observation with the pp65(NLVPMVATV):A02:01-tetramer (**Fig. 7D vs. 3B**). IE-1-specific T cells were ~51 % recipient-derived (25 recipient cells and 24 donor cells) and pp65-specific T cells were mostly recipient-derived (142 recipient cells and 20 donor cells). Taken together, these data highlight that some epitope-specific competition does appear to occur and is more stringent for pp65-specific T cells compared to IE-1-specific T cells.

### Some donor and recipient-derived clones have nearly to fully identical TCR properties

Finally, we wanted to assess how similar (in re. to TCR usage and transcriptome) newly recruited, CMV-specific T cells were compared to the recipient-derived memory population. We found 21 donor – recipient clone pairs with a statistically significant degree of TCR similarity. One of these pairs, specific for CMV IE-1, had identical TCR α and β chains including identical CDR3 regions on the amino acid level, while still differing on the nucleotide level as one would expect (**Fig. 8A**). We compared the transcriptome of these identical TCR clones and found that the recipient-derived clones had more of an effector-like phenotype (granulysin^hi^, NKG7^hi^, but CD27^lo^; **Fig. 8A**). Of note, as a trend, donor-derived T cells were expanding, while recipient-derived T cells appeared stable in abundance. Thus, a comparison at the TCR level reveals transcriptional heterogeneity and differences that were not necessarily apparent when examining all expanding clones (**Fig. 6A**). These transcriptional differences appeared to be independent of the TCR, but instead depended on the T cell’s differentiation status. Finally, patient 4 had 8 donor – recipient clone pairs specific for HLA-B*35:01-IPSINVHHY. The donor-derived clones surpassed the recipient derived clones in abundance by day 90 in all of these pairs, indicating that the donor-derived clones have an advantage that is likely to be independent of the TCR.

**Figure 8.**
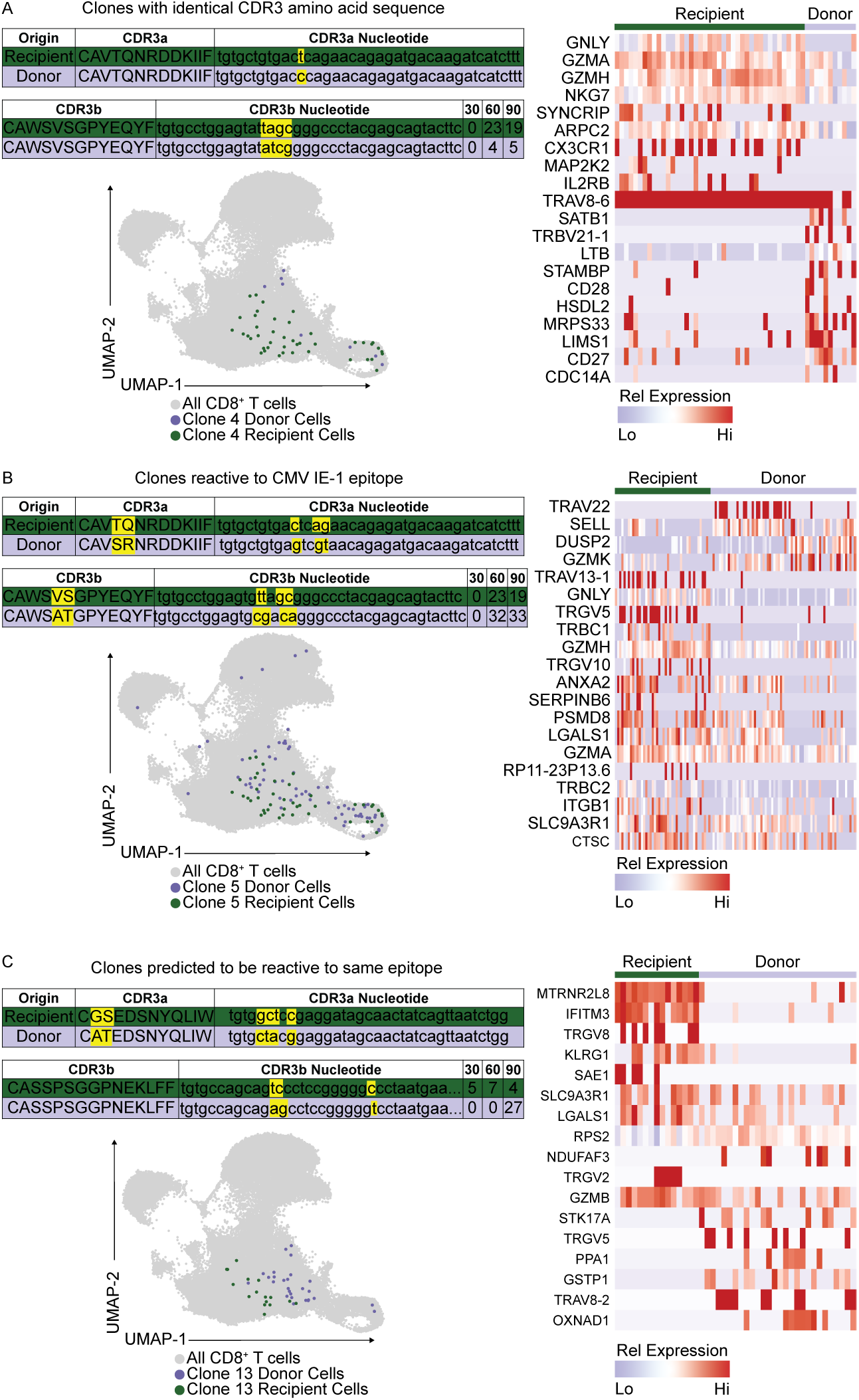
Donor- and recipient-derived T cells with identical or highly similar TCRs. (A) Shown are the CDR3 amino acid and nucleotide sequences for a donor- and recipient-derived IE-1-specific T cell clone that expanded over time with the following TCR gene usage: TRAV8-6*01, TRAJ30*01, TRBV30*01, TRBJ2-7*01. The transcriptome of each detected clone is shown in a UMAP projection. Differentially expressed genes between donor- and recipient-derived clones are highlighted in the heatmap. (B) Shown are the CDR3 amino acid and nucleotide sequences for an IE-1 specific donor- and recipient-derived T cell clone that expanded over time with the following TCR gene usage: TRAV8-6*01, TRAJ30*01, TRBV30*01, TRBJ2-7*01. The transcriptome of each detected clone is shown in a UMAP projection. Differentially expressed genes between donor- and recipient-derived clones are highlighted in the heatmap. (C) Shown are the CDR3 amino acid and nucleotide sequences for a donor- and recipient-derived T cell clone with predicted shared antigen-specificity that expanded over time with the following TCR gene usage: TRAV17*01, TRAJ33*01, TRBV28*01, TRBJ1-4*01. The transcriptome of each detected clone is shown in a UMAP projection. Differentially expressed genes between donor- and recipient-derived clones are highlighted in the heatmap.

## Discussion

We analyzed clonal T cell expansion in a unique set of longitudinal PBMC samples from patients treated with minimal myeloablative conditioning and HSCT. Following nmHSCT, patients harbored two populations of genetically unique T cells, derived from the recipient or donor. In our cohort, all recipients were CMV seropositive and hence the recipient-derived T cell compartment contained CMV-specific memory T cells. In contrast, all donors were CMV seronegative and we thus considered the donor-derived T cell compartment to be antigen-naïve. This notion is also supported a lack of donor-derived CMV-specific clones until after CMV reactivation occurred. Of note, donor and recipient pairs were not fully HLA-matched, but all donors and recipients expressed the HLA-A*02:01 allele. This is important, because it allowed us to interpret our data in context of HLA-A*02:01-restricted, naïve and memory competing T cell responses. Overall, we found that lymphocyte numbers were typically stable for the first 90 days following nmHSCT for 3 of the 4 patients indicating that homeostatic expansion during that time period was not a major driver of T cell proliferation in these patients. Furthermore, the donor-derived T cells were the predominant population in the overall CD8+ T cell compartment.

In an effort to reveal the overall clonal dynamics of the CD8+ T cell compartment, we first assessed when and how many CD8+ T cells expanded. We used a rather stringent definition of expansion which was defined as a 2-fold increase of a clone between 2 time points with at least 10 cells at any time point. This analysis revealed that expanding T cells were predominantly of donor origin and arose specifically following CMV reactivation. Although we did observe clonal expansion of recipient-derived T cells, donor-derived T cell clones expanded more vigorously. If we assume that each donor-derived clone started from a single naïve progenitor (as opposed to a small population of memory-like, homeostatically expanded T cells), then each clone would undergo approximately 24 rounds of cell division. The number of cell divisions also aligns with studies evaluating T cell expansion in mice, where a brief antigen encounter is sufficient to induce 7-10 rounds of cell division by naïve CD8+ T cells (34, 35). Prolonging the duration of TCR stimulation beyond a brief activating encounter will drive additional rounds of cell division for CD8+ T cells (34). Further mouse studies indicate that a single naïve antigen-specific T cell can generate an effector population of ~10^4^ T cells in context of an acute bacterial infection, indicating at least 14 rounds of cell division (36). This robust clonal expansion of donor-derived T cells following CMV reactivation indicates that effective clonal expansion still occurred despite the presence of recipient-derived CMV-specific memory T cells.

To specifically identify CMV-specific T cell responses in the CD8 T cell compartment, we used a combination of previously published TCR sequences and, for patient 3, ex vivo stimulation with pp65 and IE-1 peptide pools. We focused on these immunodominant epitopes to ensure that we could reliably detect antigen-specific T cells despite the inherent numeric limitations of sampling the T cell compartment with currently available single-cell sequencing based approaches (that allow for sequencing of ~10^3^-10^4^ T cells per run). While the overall CD8+ T cell compartment consisted of ~50-75% donor-derived T cells (from CMV seronegative donors), strikingly the pp65-specific T cell response was heavily dominated by recipient-derived memory T cells. However, the efficiency of competition appeared to be epitope-dependent as we observed a more pronounced donor contribution to the IE-1-specific T cell population compared to the pp65-specific T cell population - the IE-1 specific T cell response had essentially equal contributions of donor and recipient-derived T cells. These data highlight that an existing memory T cell population does not prevent *de novo* T cell responses of the same specificity. CMV-specific donor-derived T cells appear to have a competitive advantage over recipient-derived T cells, which could be related to the more terminally differentiated phenotype of the recipient-derived T cells. However, we cannot formally rule out that the condition regimen and/or graft versus host disease prophylaxis treatment (outlined in the Methods section) affected T cell function or clonal selection. Potential treatment-mediated effects are inherent confounders for which we cannot control in our study.

Remarkably, some of the newly recruited donor-derived T cells were highly similar to CMV-specific recipient clones. When comparing all donor- vs. recipient-derived clones, we observed additional instances of stunning TCR similarity, including a donor- and recipient-derived pair with fully identical TCR sequences on the amino acid level (but still showing differences on the nucleotide level). Together, these data suggest that the selection process that allows for T cell expansion is highly reproducible in humans, similar to previous observations in a mouse model system (37). Of note, the donor-derived T cells were the more abundant clones in most of the highly similar, 21 donor – recipient clone pairs indicating that the donor-derived T cells have a competitive advantage that is unrelated to TCR specificity. Transcriptome analysis of these donor-derived clones indicates that they may be more sensitive to co-stimulation (CD27) and less terminally differentiated (KLRG1, NKG2C), which may provide the observed competitive advantage (38, 39).

Finally, clonal expansion coincided with CMV reactivation, but it is possible that some of the donor-derived expanding clones were alloreactive and not CMV-specific. In nmHSCT patients, donor-derived alloresponses are essential to eliminate recipient-derived blood malignancies, but can also cause graft versus host disease (GvHD). We attempted to determine if any expanding donor-derived clones had distinct transcriptional profiles that could potentially help to discern allo-from CMV-specific responses, but we could not detect any signatures to potentially delineate these responses.

Overall, our study shows that CMV reactivation is sufficient to elicit strong *de novo* T cell responses despite the presence of CMV-specific memory T cells. We furthermore observed remarkable T cell receptor similarity between clonally expanded donor- and recipient-derived T cells suggesting reproducible selection of T cell clones with congruent T cell receptor specificities, which overall appears to lead to a rejuvenation of the memory T cell pool without a pronounced change in the TCR repertoire.

## Acknowledgements and funding sources

We thank Nicholas J. Maurice for critical review of the manuscript, and David Levine for assistance with genotyping patient 3. This work was supported by National Institutes of Health (NIH) grants R01AI123323 (to M.P.). J.R.E. was supported by NIH T32 AI007509. P.B. was supported by R01AI136514 and ORIP S10OD028685 to support Fred Hutch Scientific Computing. The sample repository was supported by the Fred Hutch Vaccine and Infectious Disease Division.

## Materials and Methods

### Human Cohort and sample processing

This study was approved by the Fred Hutchinson Cancer Research IRB and all subjects signed informed consent. The patient cohort contained four recipient - donor pairs with CMV reactivation events between day 30 and 90 after transplantation each with the following clinical diagnosis prior to nmHSCT: Patient 1 was diagnosed with Myelodysplasia (recipient 68 years, donor 38 years; this was the patient’s 2^nd^ HSCT procedure); Patient 2 was diagnosed with Mantle Cell Lymphoma (recipient 60 years, donor 29 years); Patient 3 was diagnosed with acute lymphocytic leukemia (recipient 57 years, donor 55 years); Patient 4 was diagnosed with non-Hodgkin’s lymphoma (recipient 60 years, donor 20 years). Patients received fludarabine and 200-300 cGy of total body irradiation for pretransplant conditioning. Peripheral blood stem cells served as the graft source. Patients 2, 3 and 4 received cyclosporine and mycophenolate mofetil (MMF) for GVH prophylaxis. Patient 1 was treated with sirolimus in addition to cyclosporine and MMF. CMV surveillance was done weekly by PCR and patients were preemptively treated with (val)ganciclovir as recently described (24). Peripheral blood mononuclear cells (PBMCs) from each patient and for each time point were obtained as cryopreserved samples from the Infectious Disease Sciences Biospecimen Repository, Vaccine and Infectious Disease Division, FHCRC. Vials with cryopreserved cells were thawed at 37°C until a tiny ice crystal was left in the tube, and then carefully diluted in 1mL of pre-warmed complete RPMI (RPMI (Gibco, #18875119) with 10% FBS (Nucleus Biologics, #AU FBS-500ml L1 HI) and 1% Penicillin-Streptomycin (Gibco, #15140122) and transferred to a new tube. An additional 13 mL of pre-warmed complete RPMI were added drop by drop, followed by centrifugation for 5 minutes at 400g and resuspension in 1 mL of complete RPMI.

### T cell stimulation assay

Freshly thawed PBMCs were resuspended at 10^7^ cells/mL. 100μL of cells were added into a 96-well round bottom plate. 50 μL stimulation cocktail was added to each well containing cells. Stimulation cocktail was made by adding 1μg/mL of both anti-CD28 and anti-CD49d (BD Biosciences) with 2 μg/mL of either overlapping peptide pools of pp65, IE-1 (pp65 and IE-1 PepMix, JPT peptide technologies), or DMSO into complete RPMI. PBMCs with stimulation cocktail was incubated at 37°C for 18 hours. Cells were then prepared for FACS.

### Flow Cytometry and Cell sorting

For flow cytometric analysis, good practices were followed as outlined in the guidelines for use of flow cytometry (40). Following thawing or stimulation, PBMCs were incubated with Fc-blocking reagent (BioLegend Trustain FcX, #422302) and fixable Aqua Live/Dead reagent (ThermoFisher, #L34957) in PBS (Gibco, #14190250) for 15 minutes at room temperature. If required, cells were stained with an CMV-Tetramer reagent (peptide NLVPMVATV; NIH Tetramer Core) diluted in FACS buffer (PBS with 2% FBS, Nucleus Biologics, #AU FBS-500ml L1 HI) for 30 minutes at room temperature, followed by two washes. After this, cells were incubated for 20 minutes at room temperature with 100 μl total volume of antibody master mix freshly prepared in Brilliant staining buffer (BD Bioscience, #563794), followed by two washes. All antibodies were titrated and used at optimal dilution, and staining procedures were performed in 96-well round-bottom plates. Stained PBMCs were resuspended in FACS buffer and sorted.

All cell sorting was performed on a FACSAria III (BD Biosciences), equipped with 20 detectors and 405nm, 488nm, 532nm and 628nm lasers. For all sorts, an 85 μm nozzle operated at 45 psi sheath pressure was used. Single-stained controls were prepared with every experiment using antibody capture beads diluted in FACS buffer (BD Biosciences anti-mouse, #552843 and anti-rat, #552844), or cells for Live/Dead reagent. Cells were sorted into chilled Eppendorf tubes containing 500 μL of complete RPMI, washed once in PBS and immediately used for subsequent processing.

### Single-cell library preparation and sequencing

cDNA libraries of CMV-Tetramer+ CD8+ T cells were generated using the Chromium Single Cell 3’ Reagent Kits v2 while CMV-Tetramer-CD8+ T cells and CD8+ stimulated T cells were generated using the Chromium Single Cell 5’ Reagent Kits v1 with Human T cell V(D)J enrichment kits (10x Genomics). The Chromium Single Cell protocol targeting 10,000 cells per well was followed. Briefly, single cells were isolated into oil emulsion droplets with barcoded gel beads and reverse transcriptase mix. cDNA was generated within these droplets, then the droplets were dissociated. cDNA was purified using DynaBeads MyOne Silane magnetic beads (ThermoFisher, #370002D). cDNA amplification was performed by PCR (10 cycles) using reagents within the Chromium Single Cell 3’ Reagent Kit v2 (10x Genomics). Amplified cDNA was purified using SPRIselect magnetic beads (Beckman Coulter). If necessary, target enrichment was also performed by PCR (10 cycles) and cDNA purification via SPRISelect beads. cDNA was enzymatically fragmented and size selected prior to library construction. Libraries were constructed by performing end repair, A-tailing, adaptor ligation, and PCR (12 cycles). Quality of the libraries was assessed by using Agilent 2200 TapeStation with High Sensitivity D5000 ScreenTape (Agilent). Quantity of libraries was assessed by performing digital droplet PCR (ddPCR) with Library Quantification Kit for Illumina TruSeq (BioRad, #1863040). Libraries were diluted to 2 nM and paired-end sequencing was performed on a HiSeq 2500 sequencer (Illumina). Stimulation libraries were diluted to 3nM and paired-end sequencing was performed on a NovaSeq 6000 (Illumina).

### Sequencing Data Processing

Raw base call (BCL) files were demultiplexed to generate Fastq files using the cellranger mkfastq pipeline within Cell Ranger 2.1.1 (10x Genomics). Targeted transcriptome Fastqs were further analyzed via Seven Bridges (BD Biosciences). Whole transcriptome Fastq files were processed using the standard cellranger pipeline (10x genomics) within Cell Ranger 2.1.1. Briefly, cellranger count performs alignment, filtering, barcode counting, and UMI counting. The cellranger count output was fed into the cellranger aggr pipeline to normalize sequencing depth between samples. The final output of cellranger (molecule per cell matrix) was then analyzed in R using the package Seurat (version 2.3 and 3.0) as described below.

### Sequencing analysis

The R package Seurat (41) was utilized for all downstream analysis. For whole transcriptome data, based on commonly used cutoffs suggested by Butler et al, only cells that had at least 200 genes (with ≤ 20% being mitochondrial genes) were included in analysis (removing 182 out of a total of 5,416 cells). A natural log normalization using a scale factor of 10,000 was performed across the library for each cell. UMIs and mitochondrial genes were linearly scaled to remove these variables as unwanted sources of variation. Doublets and low-quality cells were identified by their outlier UMI and gene counts on a per patient basis, and their high percentage of mitochondrial genes (more than 20%).

For whole transcriptome analysis (WTA), dimensionality reduction using UMAP and clustering was performed on a subset of variable genes. When scaling data, UMI was the only regressed variable. Dimensionality reduction using UMAP and clustering was based on either all genes or all proteins. For differential gene expression analysis we utilized the Seurat implementation of MAST (model-based analysis of single-cell transcriptomes) with the number of UMIs included as a covariate (proxy for cellular detection rate (CDR)) in the model (42). To combine datasets, Harmony was used (43).

Genotype-informative SNPs in single-cell transcripts were identified by correlation analysis of heterozygous positions across cells followed by clustering to define groups of covarying SNPs. Sex-specific gene expression (patients 2 and 4) and previous genotyping data (patient 3) were then used to disambiguate donor and recipient. Genotype calls at the single-cell level were compared with output from the Sourporcell algorithm (44) and found to be greater than 99% concordant. TCR sequence matching was performed using the TCRdist algorithm as implemented in the Clonotype Neighbor Graph Analysis (CoNGA) python package’s find_significant_tcrdist_matches function (https://github.com/phbradley/conga) (45). In this approach, the TCRdist score for a match is compared to a background distribution of TCRdist scores for the same TCRs matched to random TCR sequences generated using a probabilistic model of the V(D)J recombination process.

## DATA AND CODE AVAILABILITY

The sequencing data discussed in this publication have been deposited in the NCBI’s Gene Expression Omnibus (46) and are accessible through GEO series accession number GSE167825. (https://www.ncbi.nlm.nih.gob/geo/query/acc/cgi?acc=GSE167825). All scripts used for data processing and plot generation are available at https://github.com/Jami-Erickson/scRNAseq_CD8Tcells_CMV.

---

**Supplemental Figure 1.**
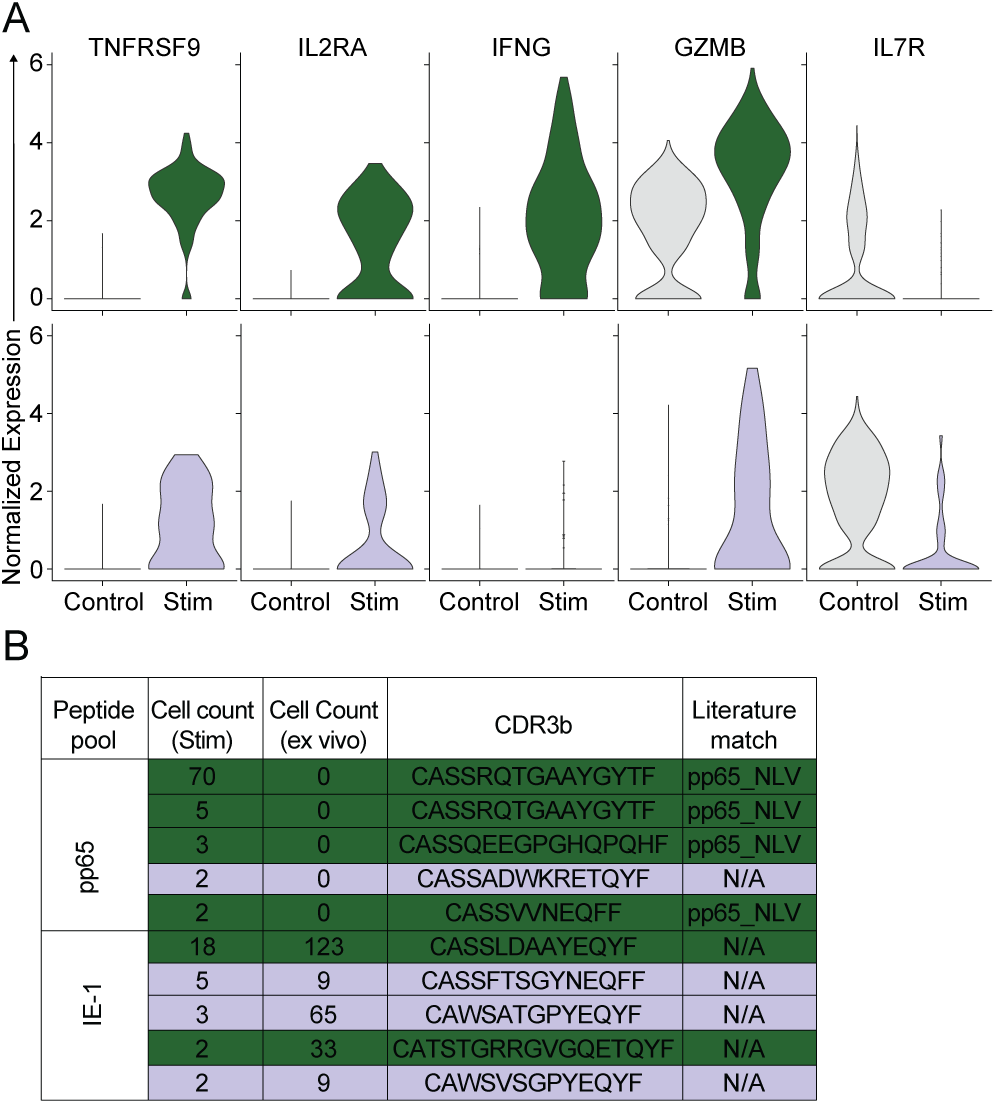
Additional analysis of peptide-pool stimulated CD137+ CD8+ T cells compared to unstimulated CD8 T cells (control). (A) The gene expression profile of recipient-derived (green) and donor-derived (purple) CD8 T cells compared to CD8 T cells from the DMSO (no stim) control group (grey). (B) CD137+ sorted T cells were analyzed by 5’ scRNA-seq to determine TCR gene usage and CDR3 sequences. Cell counts for each clone are shown from the peptide pool stimulation experiment (stim) and compared to the direct ex vivo sequencing data (shown in Fig. 3).

## References

1. Qi Q, et al. (2014) Diversity and clonal selection in the human T-cell repertoire. Proc Natl Acad Sci U S A 111(36):13139–13144.

2. Liacopoulos P & Ben-Efraim S (1975) Antigenic competition. Prog Allergy 18:97–204.

3. Kedl RM, Kappler JW, & Marrack P (2003) Epitope dominance, competition and T cell affinity maturation. Curr Opin Immunol 15(1):120–127.

4. Oberle SG, et al. (2016) A Minimum Epitope Overlap between Infections Strongly Narrows the Emerging T Cell Repertoire. Cell Rep 17(3):627–635.

5. Johnson LR, Weizman OE, Rapp M, Way SS, & Sun JC (2016) Epitope-Specific Vaccination Limits Clonal Expansion of Heterologous Naive T Cells during Viral Challenge. Cell Rep 17(3):636–644.

6. den Haan JM, et al. (1995) Identification of a graft versus host disease-associated human minor histocompatibility antigen. Science 268(5216):1476–1480.

7. Wallny HJ & Rammensee HG (1990) Identification of classical minor histocompatibility antigen as cell-derived peptide. Nature 343(6255):275–278.

8. Grufman P, Wolpert EZ, Sandberg JK, & Kärre K (1999) T cell competition for the antigen-presenting cell as a model for immunodominance in the cytotoxic T lymphocyte response against minor histocompatibility antigens. Eur J Immunol 29(7):2197–2204.

9. Kedl RM, et al. (2000) T cells compete for access to antigen-bearing antigen-presenting cells. J Exp Med 192(8):1105–1113.

10. Zehn D, Roepke S, Weakly K, Bevan MJ, & Prlic M (2014) Inflammation and TCR signal strength determine the breadth of the T cell response in a bim-dependent manner. J Immunol 192(1):200–205.

11. Badovinac VP, Haring JS, & Harty JT (2007) Initial T cell receptor transgenic cell precursor frequency dictates critical aspects of the CD8(+) T cell response to infection. Immunity 26(6):827–841.

12. Butz EA & Bevan MJ (1998) Massive expansion of antigen-specific CD8+ T cells during an acute virus infection. Immunity 8(2):167–175.

13. Frahm N, et al. (2012) Human adenovirus-specific T cells modulate HIV-specific T cell responses to an Ad5-vectored HIV-1 vaccine. J Clin Invest 122(1):359–367.

14. Borysiewicz LK, Morris S, Page JD, & Sissons JG (1983) Human cytomegalovirus-specific cytotoxic T lymphocytes: requirements for in vitro generation and specificity. Eur J Immunol 13(10):804–809.

15. McLaughlin-Taylor E, et al. (1994) Identification of the major late human cytomegalovirus matrix protein pp65 as a target antigen for CD8+ virus-specific cytotoxic T lymphocytes. J Med Virol 43(1):103–110.

16. Walter EA, et al. (1995) Reconstitution of cellular immunity against cytomegalovirus in recipients of allogeneic bone marrow by transfer of T-cell clones from the donor. N Engl J Med 333(16):1038–1044.

17. Khan N, Cobbold M, Keenan R, & Moss PA (2002) Comparative analysis of CD8+ T cell responses against human cytomegalovirus proteins pp65 and immediate early 1 shows similarities in precursor frequency, oligoclonality, and phenotype. J Infect Dis 185(8):1025–1034.

18. Sylwester AW, et al. (2005) Broadly targeted human cytomegalovirus-specific CD4+ and CD8+ T cells dominate the memory compartments of exposed subjects. J Exp Med 202(5):673–685.

19. Jackson SE, Mason GM, Okecha G, Sissons JG, & Wills MR (2014) Diverse specificities, phenotypes, and antiviral activities of cytomegalovirus-specific CD8+ T cells. J Virol 88(18):10894–10908.

20. McSweeney PA, et al. (2001) Hematopoietic cell transplantation in older patients with hematologic malignancies: replacing high-dose cytotoxic therapy with graft-versus-tumor effects. Blood 97(11):3390–3400.

21. Slavin S, et al. (1998) Nonmyeloablative stem cell transplantation and cell therapy as an alternative to conventional bone marrow transplantation with lethal cytoreduction for the treatment of malignant and nonmalignant hematologic diseases. Blood 91(3):756–763.

22. Gyurkocza B & Sandmaier BM (2014) Conditioning regimens for hematopoietic cell transplantation: one size does not fit all. Blood 124(3):344–353.

23. Cannon MJ, Schmid DS, & Hyde TB (2010) Review of cytomegalovirus seroprevalence and demographic characteristics associated with infection. Rev Med Virol 20(4):202–213.

24. Zamora D, et al. (2021) Cytomegalovirus-specific T-cell reconstitution following letermovir prophylaxis after hematopoietic cell transplantation. Blood 138(1):34–43.

25. Dash P, et al. (2017) Quantifiable predictive features define epitope-specific T cell receptor repertoires. Nature 547(7661):89–93.

26. Islam S, et al. (2011) Characterization of the single-cell transcriptional landscape by highly multiplex RNA-seq. Genome Res 21(7):1160–1167.

27. Ramsköld D, et al. (2012) Full-length mRNA-Seq from single-cell levels of RNA and individual circulating tumor cells. Nat Biotechnol 30(8):777–782.

28. Hashimshony T, Wagner F, Sher N, & Yanai I (2012) CEL-Seq: single-cell RNA-Seq by multiplexed linear amplification. Cell Rep 2(3):666–673.

29. Martin PJ, et al. (2020) Recipient and donor genetic variants associated with mortality after allogeneic hematopoietic cell transplantation. Blood Adv 4(14):3224–3233.

30. Becht E, et al. (2018) Dimensionality reduction for visualizing single-cell data using UMAP. Nat Biotechnol.

31. Aran D, et al. (2019) Reference-based analysis of lung single-cell sequencing reveals a transitional profibrotic macrophage. Nat Immunol 20(2):163–172.

32. Shugay M, et al. (2018) VDJdb: a curated database of T-cell receptor sequences with known antigen specificity. Nucleic Acids Res 46(D1):D419–D427.

33. Wolfl M, et al. (2007) Activation-induced expression of CD137 permits detection, isolation, and expansion of the full repertoire of CD8+ T cells responding to antigen without requiring knowledge of epitope specificities. Blood 110(1):201–210.

34. Prlic M, Hernandez-Hoyos G, & Bevan MJ (2006) Duration of the initial TCR stimulus controls the magnitude but not functionality of the CD8+ T cell response. J Exp Med 203(9):2135–2143.

35. Kaech SM & Ahmed R (2001) Memory CD8+ T cell differentiation: initial antigen encounter triggers a developmental program in naive cells. Nat Immunol 2(5):415–422.

36. Neuenhahn M, et al. (2006) CD8alpha+ dendritic cells are required for efficient entry of Listeria monocytogenes into the spleen. Immunity 25(4):619–630.

37. Cukalac T, et al. (2014) Reproducible selection of high avidity CD8+ T-cell clones following secondary acute virus infection. Proc Natl Acad Sci U S A 111(4):1485–1490.

38. Ibegbu CC, et al. (2005) Expression of killer cell lectin-like receptor G1 on antigen-specific human CD8+ T lymphocytes during active, latent, and resolved infection and its relation with CD57. J Immunol 174(10):6088–6094.

39. Prlic M, Sacks JA, & Bevan MJ (2012) Dissociating markers of senescence and protective ability in memory T cells. PLoS One 7(3):e32576.

40. Cossarizza A, et al. (2019) Guidelines for the use of flow cytometry and cell sorting in immunological studies (second edition). Eur J Immunol 49(10):1457–1973.

41. Butler A, Hoffman P, Smibert P, Papalexi E, & Satija R (2018) Integrating single-cell transcriptomic data across different conditions, technologies, and species. Nature biotechnology 36(5):411–420.

42. Finak G, et al. (2015) MAST: a flexible statistical framework for assessing transcriptional changes and characterizing heterogeneity in single-cell RNA sequencing data. Genome biology 16(1):278.

43. Korsunsky I, et al. (2019) Fast, sensitive and accurate integration of single-cell data with Harmony. Nat Methods 16(12):1289–1296.

44. Heaton H, et al. (2020) Souporcell: robust clustering of single-cell RNA-seq data by genotype without reference genotypes. Nat Methods 17(6):615–620.

45. Schattgen SA, et al. (2021) Integrating T cell receptor sequences and transcriptional profiles by clonotype neighbor graph analysis (CoNGA). Nat Biotechnol.

46. Edgar R, Domrachev M, & Lash AE (2002) Gene Expression Omnibus: NCBI gene expression and hybridization array data repository. Nucleic Acids Res 30(1):207–210.

